# BnIR: a multi-omics database with various tools for *Brassica napus* research and breeding

**DOI:** 10.1101/2023.01.12.523736

**Authors:** Zhiquan Yang, Shengbo Wang, Lulu Wei, Yiming Huang, Dongxu Liu, Yupeng Jia, Chengfang Luo, Yuchen Lin, Congyuan Liang, Yue Hu, Cheng Dai, Liang Guo, Yongming Zhou, Qing-Yong Yang

**Affiliations:** National Key Laboratory of Crop Genetic Improvement, Hubei Hongshan Laboratory, Huazhong Agricultural University, Wuhan, 430070, China; Hubei Key Laboratory of Agricultural Bioinformatics, College of Informatics, Huazhong Agricultural University, Wuhan, 430070, China; Innovative Center of Molecular Genetics and Evolution, School of Life Sciences, Guangzhou University, Guangzhou, 510405, China

**Keywords:** *Brassica napus*, multi-omics, database, candidate gene mining, functional genomics

## Abstract

In the post-GWAS era, multi-omics techniques have shown great power and potential for candidate gene mining and functional genomics research. However, due to the lack of effective data integration and multi-omics analysis platforms, such techniques have not still been applied widely in rapeseed, an important oil crop worldwide. Here, we constructed a rapeseed multi-omics database (BnIR; http://yanglab.hzau.edu.cn/BnIR), which provides datasets of six omics including genomics, transcriptomics, variomics, epigenetics, phenomics and metabolomics, as well as numerous “variation-gene expression-phenotype” associations by using multiple statistical methods. In addition, a series of multi-omics search and analysis tools are integrated to facilitate the browsing and application of these datasets. BnIR is the most comprehensive multi-omics database for rapeseed so far, and two case studies demonstrated its power to mine candidate genes associated with specific traits and analyze their potential regulatory mechanisms.

**Short Summary:** In this study, we developed BnIR (http://yanglab.hzau.edu.cn/BnIR), a multi-omics database for rapeseed that integrates six omics datasets and “variation-gene expression-phenotype” associations. The database is equipped with a range of multi-omics tools for streamlined browsing and analysis. Through multiple case studies, we demonstrated BnIR’s potential to identify candidate genes associated with specific traits and investigate their regulatory mechanisms.

## Introduction

For comprehensive understanding of the mechanism underlying crop trait formation, it is important to interpret the molecular intricacy and variation at multiple levels such as the genome, epigenome, transcriptome, proteome, metabolome and phenome (Edwards and Batley, 2004; Li et al., 2016; Fernie and Gutierrez-Marcos, 2019; Tuggle et al., 2022). In recent years, the rapid advancement in multi-omics techniques has provided researchers with more multidimensional information to dissect complex biological systems (Ichihashi et al., 2020; Wu et al., 2021; Yang et al., 2021). Based on massive multi-omics datasets, numerous research methods have been developed to mine genetic loci and candidate genes regulating phenotypes, such as genome-wide association study (GWAS) (Yu et al., 2006), expression quantitative trait locus (eQTL) mapping(Goring et al., 2007), transcriptome-wide association study (TWAS) (Gusev et al., 2016) and summary data-based Mendelian randomization analysis (SMR) (Zhu et al., 2016). These methods have high efficiencies in mining genetic loci and candidate genes associated with certain traits and have been widely applied in crop breeding (Yu *et al*., 2006; Wang et al., 2018; Lu et al., 2019; Liu et al., 2020; Tang et al., 2021; Zhang et al., 2021; He et al., 2022; Zhang et al., 2022a; Zhang et al., 2022b). Multi-omics techniques are becoming increasingly important in assisting crop breeding (Yang *et al*., 2021). For more efficient utilization of multi-omics data, multi-omics databases have been developed in some important crops, such as wheat (*Triticum aestivum* L.) (Ma et al., 2021), soybean (*Glycine max*) (Grant et al., 2010), tomato (*Solanum lycopersicum*) (Kudo et al., 2017), barley (*Hordeum vulgare* L.) (Li et al., 2022), maize (*Zea mays* L.) (Gui et al., 2020), millet (*Setaria italica* L.) (Yang et al., 2020), cotton (*Gossypium hirsutum* L.) (Yang et al., 2022b) and rice (*Oryza sativa* L.) (Sakai et al., 2013).

Rapeseed is the second most important oilseed crop widely planted in the world. The traits including yield, seed oil content (SOC), oil quality, disease resistance and stress resistance largely determine its production and economic value (Friedt et al., 2018). Due to its special origin and evolutionary history, rapeseed has a polyploid genome and therefore a more complex sequence composition and regulatory mechanism than diploid crops, which increases the difficulty in fine mapping of the candidate loci and genes (Chalhoub et al., 2014; Friedt et al., 2018). Recently, a large number of genetic loci and candidate genes associated with important traits have been efficiently identified using powerful multi-omics techniques and analysis tools (Lu *et al*., 2019; Tang *et al*., 2021; He *et al*., 2022; Hu et al., 2022; Zhang *et al*., 2022b). However, the identification is still rather time-consuming, and the regulatory mechanisms of most identified candidate loci or genes on phenotypes remain elusive. In our recent work, we have constructed and released several databases, including BnPIR (Song et al., 2021) (http://cbi.hzau.edu.cn/bnapus/), BnTIR (Liu et al., 2021a) (http://yanglab.hzau.edu.cn/BnTIR/) and BnVIR (Yang et al., 2022a) (http://yanglab.hzau.edu.cn/BnVIR/), by collecting multi-omics data including genome, transcriptome and population genetic variations and developing related omics tools. These databases have been widely applied to facilitate related research work. However, there is still a lack of platforms that integrate multi-omics data to help genetic breeding and functional genomics analysis in rapeseed.

In this study, we constructed BnIR (http://yanglab.hzau.edu.cn/BnIR/), the first *Brassica napus* multi-omics information resource database. BnIR integrates the most comprehensive multi-omics datasets involving various dimensions. In the database, multiple association analysis tools can be applied to mine the genetic loci, candidate genes and genetic variations, providing important resources, tools and references for rapeseed breeding. In addition, BnIR provides numerous common online multi-omics data analysis tools, which can support all published rapeseed genomes and offers a convenient and efficient analysis platform. We believe that BnIR will be a valuable database for future rapeseed breeding and functional genomics research.

## Results

### Construction and overview of BnIR

For a comprehensive understanding of rapeseed genomics, we first acquired and integrated rapeseed multi-omics data from recent research and databases. In total, datasets from six omics were obtained, including 29 genome assemblies (Supplementary Table S1), genetic variations of 2,311 accessions, transcriptome data from 2,791 libraries (Supplementary Table S2), 118 phenotypes (Supplementary Table S3), the contents of 266 metabolites and epigenome signals involving DNA methylation, histone modification, chromatin accessibility and chromatin interaction (Supplementary Table S4). Next, the related information was mined by processing and analyzing of these datasets. In addition, multiple multi-omics analysis methods were employed to mine the genetic loci, candidate genes and genetic variations, including GWAS, eQTL, TWAS, SMR and co-localization analysis. Finally, a comprehensive rapeseed multi-omics information resource database (BnIR; http://yanglab.hzau.edu.cn/BnIR/) was constructed (Figure 1).

**Figure 1.**
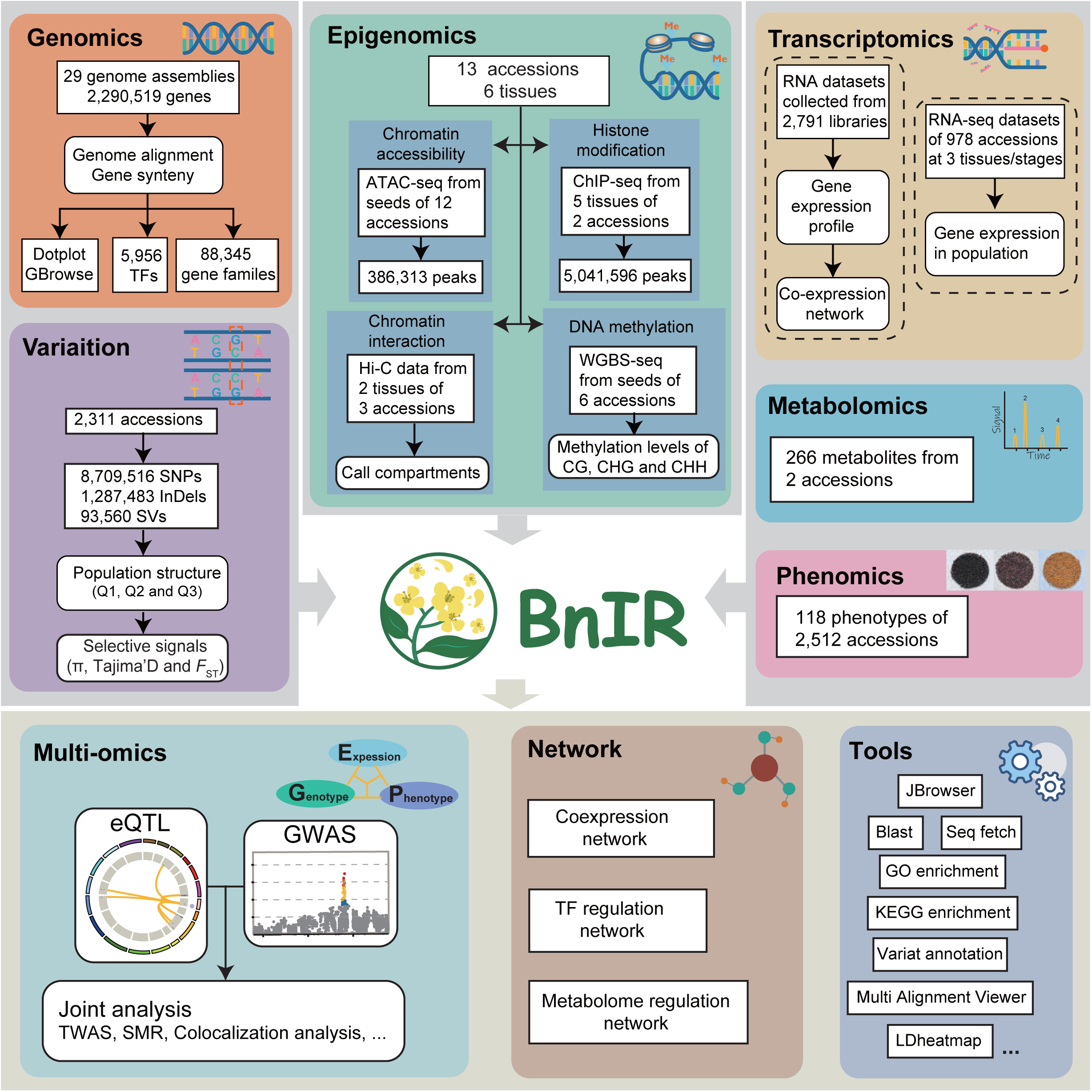
Construction pipelines and overview of BnIR.

BnIR comprises 12 portals, including Population, Genomics, Transcriptomics, Variation, Phenotype, Epigenetics, Metabolome, Multi-omics and Network, Tools, Download and Help. Abundant and convenient visual tools are provided in these portals for browsing and comparing the germplasm resource information, genome sequences, gene structures, epigenetic signals, metabolite contents and phenotypes, so as to manage germplasm resource and assist the exploring of mechanism underlying gene regulation and evolution.

### New features in omics portals

#### Genomics

Based on our previous BnPIR database, we added Darmor (v.10), Ningyou7 and Express617 genome assemblies as well as their genome sequences and annotations, which can be downloaded from the Genome data module (Supplementary Table S1). Genome synteny and gene index have been updated, which can be browsed in the Gbrowse synteny and Gene index modules. According to the phylogenetic trees of homologous genes, the Gene search module was developed for users to browse and compare the gene structures to determine whether the homologous genes have undergone functional differentiation at the sequence level. In addition, the gene families and transcription factors (TF) of 12 *B. napus* genome assemblies are annotated, which can be browsed in the corresponding modules of the Genomics portal.

#### Transcriptomics

As for the Transcriptomics portal, the previous core function modules in BnTIR, including gene expression pattern query and eFP browser, have been integrated into eFP(single gene module) and eFP(multiple gene module) of BnIR. In two eFP modules, the webpage has been redesigned to greatly improve the response speed by more than 150 times (from 9.348 s to 0.0611 s). In addition, the published RNA datasets from 537 RNA-seq libraries involving seven hormone treatments and six abiotic stress treatments have been added into the eFP modules (Supplementary Table S2). Users can browse and compare the gene expression pattern between the treatment and control groups.

In order to acquire gene expression data from more accessions, the RNA-seq data of 2,791 published tissue samples were collected, processed and summarized (Supplementary Table S2). Finally, the gene expression data of these samples were integrated into the Expression profile (meta library) module of the Transcriptomics portal, in which users can browse the expression of interested genes in all samples. As for the breeding of *B. napus*, ecotype and seed quality improvement are two key periods. In addition, heterosis is an important phenomenon and widely used for genetic improvement. To facilitate studies of the genes involved in these improvement processes, we collected the gene expression datasets from eight representative cultivars of three ecotypes, double-high (high erucic acid and high glucosinolate) and double-low (low erucic acid and low glucosinolate) cultivars (Zhongyou821 and Zhongshuang11), and eight RNA-seq datasets of accessions with heterosis. Users can browse and compare the expression levels of interested genes between different accessions in the Expression profile (meta library) module to determine whether the gene is related to the breeding improvement process. In addition, we collected and processed the population-level gene expression datasets in leaves and seeds at 20 and 40 days after flowering (DAF). Users can browse the population-level expression patterns of the interested genes in the Population expression module. In summary, new transcriptomic information and tools will help researchers to understand the function and expression features of the interested genes from a broader perspective.

#### Variation and phenotype

The previous BnVIR database integrated 10,090,561 genetic variations and 21 phenotypes of 2,311 core *B. napus* accessions (Yang et al., 2022a). On this basis, we newly collected 216 published phenotype datasets from 13 studies (Bus et al., 2011; Sun et al., 2016b; Sun et al., 2016a; Chen et al., 2018; Kittipol et al., 2019; Wu et al., 2019; Song et al., 2020; Xuan et al., 2020; Liu et al., 2021c; Liu et al., 2021b; Wang et al., 2021; Zhang *et al*., 2021; Zhang *et al*., 2022a) (Supplementary Table S3). These datasets cover a variety of phenotypic traits, such as agronomic traits under abiotic stress conditions and seed quality traits, and have been integrated into the Phenotype and Variation portals. Users can browse the phenotypic values and compare them between the accessions of different ecotypes in the Phenotype portal to check whether the phenotype is related to ecotype improvement. In the Variation portal, users can compare the phenotypic value between accessions with different alleles or haplotypes to explore the effect of genetic variation on these phenotypes (Korber et al., 2015; Korber et al., 2016)

#### Epigenome

For epigenetic data, we collected the ChIP-seq and WGBS datasets for five tissues of six accessions including six histone modifications, ATAC-seq datasets for seeds of four accessions and Hi-C datasets of three accessions (Supplementary Table S4). After filtering and processing, a total of 5,041,596 ChIP-seq peaks and 386,313 ATAC-seq peaks were obtained, which were integrated into the Histone modification and Chromatin accessibility modules, respectively. Based on the WGBS datasets, the CG, CHG and CHH methylation levels of genes were obtained and integrated into the DNA methylation module. Based on the Hi-C datasets, chromatin interaction frequency and compartments were calculated and integrated into the Chromatin interaction module. In addition, users can browse the tracks of all epigenetic data of the gene segment of interest using the JBrowser or query these epigenetic datasets in the corresponding module of the Epigenetic portal. These datasets integrated the epigenetic information involving histone modification, epigenomic state, chromatin accessibility and interaction, providing the first systematic epigenetic data resource in *B. napus* to help identify regulatory elements in *B. napus* genome and understand their regulatory mechanism.

#### Metabolome

In the Metabolome portal, the data about the contents of 266 metabolites of two accessions from two studies were collected and integrated. Users can query the related information of these metabolites, such as structure, molecular formula and content in these accessions in the corresponding module, which provides a way to quickly obtain the information of metabolites.

### Integration and analysis of multi-omics data

Based on the collected population-level multi-omics data, we performed GWAS, eQTL and TWAS to mine candidate genes and genetic variations associated with phenotypes. A total of 379 loci were identified to be significantly associated with 55 phenotypes by GWAS, involving 2,119 candidate genes (Supplementary Table S5). By using the population-level gene expression datasets from seeds at 20 and 40 DAF of 309 accessions, 1,118,218 eQTLs involving 968,375 lead SNPs (eSNPs) and 33,507 genes (eGenes) were identified, including 356,112 *cis*-eQTLs and 762,106 *trans*-eQTLs. In total, 3,724 genes were identified to be significantly associated with 124 phenotypes by TWAS (Supplementary Table S7). Next, to more precisely identify the candidate genes associated with phenotypes by combining the three omics datasets, co-localization and SMR analyses were performed based on the GWAS and eQTL results (Supplementary Tables S8 and S9). As a result, co-localization analysis identified 130 genes significantly associated with 24 phenotypic datasets; SMR identified 298 genes significantly associated with 45 phenotypic datasets. By combining TWAS, co-localization analysis and SMR results, a total of 3,991 genes were found to be significantly associated with 131 phenotypes (Figure 2A). Known associations reported in previous studies were also identified in our analysis, such as *BnaC09.MYB28* for seed glucosinolate content (SGC) (Case study 1), and *BnaA05.PMT6* for SOC, suggesting that the analysis results are reproducible (Supplementary Table S8). In addition, some new candidate genes were found to be related to important traits, such as *BnaC07.SRK2D* for SOC (Case study 2), suggesting that the analysis can be used to mine new candidate genes.

**Figure 2.**
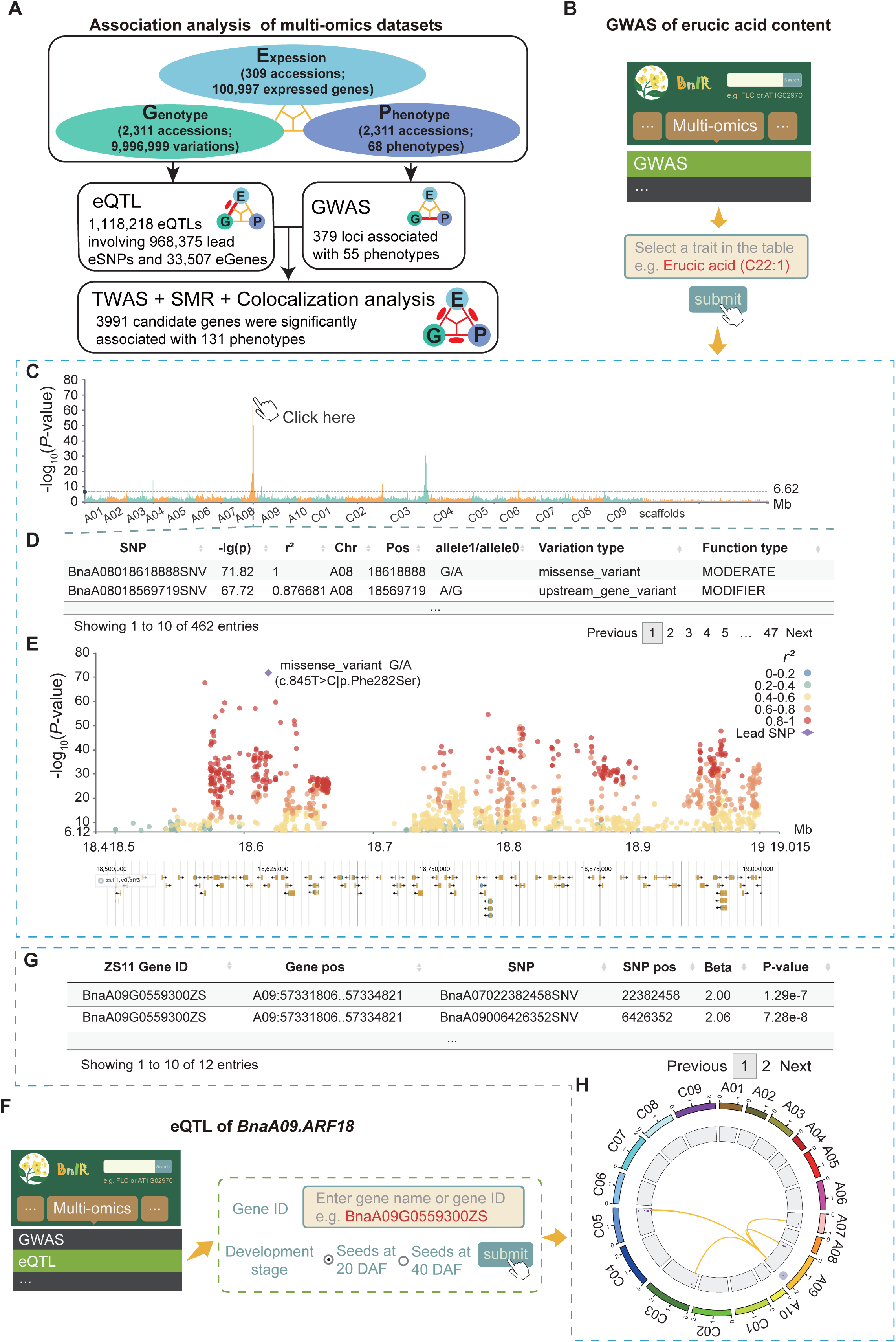
Integration of multi-omics data. **(A)** Pipeline of integrating population-level multi-omics datasets. **(B-E)** Usage of the GWAS module. **(B)** Pipeline of searching the GWAS signals of seed erucic acid content. **(C)** Global GWAS signals of seed erucic acid content. **(D)** List of significant variations in A08:18500000–19000000. **(E)** Local Manhattan plot of A08:18500000-19000000. Purple diamond represents the lead variation. Green to red dots indicate weak to strong linkage disequilibrium coefficients with the lead variation. **(F-H)** Usage of eQTL module. **(F)** Pipeline of searching the eQTL signals of *BnaA09.ARF18*. **(G)** List of eSNPs of *BnaA09.ARF18*. **(H)** Circos plot of the global eQTLs of *BnaA09.ARF18*. Purple dot in the outer circle represents the genomic position of *BnaA09.ARF18*. Purple dots in the median circle represent the P- values of eSNPs. Yellow lines represent the associations between eSNPs and *BnaA09.ARF18*.

Based on the above results, GWAS, eQTL, TWAS, COLOC, and SMR modules in the Multi-omics portal were designed for convenient browsing and use of these datasets. For example, users can browse information about the global GWAS signals and significant variations for the phenotypes of interest in the GWAS module. Taking the erucic acid content as an example, when users select 8Erucic acid9 in the Trait box and submit it (Figure 2B), the page will display the Manhattan plot of global GWAS results, showing that the locus on chromosome A08 is the most significant locus (Figure 2C). Then, the user can scroll the mouse to magnify this locus and select the region of <A08:18500000–19000000= to browse for the related information of significant variations (Figure 2D). Combined with the gene annotations, two non-synonymous mutations in the reported candidate gene (*BnaA08.FAE1*) for erucic acid content in seeds were both identified in our GWAS results (Figure 2E). In the eQTL module, users can browse the genome-wide information about *cis*-eSNPs and *trans*-eSNPs regulating gene expression levels. For example, the expression level of *BnaA09.ARF18* was reported to be associated with the silique length and thousand-seed weight (Liu et al., 2015). When users enter 8BnaA09G0559300ZS9 (*BnaA09.ARF18*) and submit it (Figure 2F), the page will display a list and a circus figure of 12 eQTLs regulating its expression level (Figure 2G), including four *cis*-eSNPs near *BnaA09.ARF18* and eight *trans*-eSNPs from four chromosomes (Figure 2H). Additionally, users can browse the information of candidate genes associated with phenotypes identified by three methods respectively in the TWAS, SMR, and COLOC modules.

### Development and application of multi-omics tools

To provide a more convenient and user-friendly platform for researchers, we developed and integrated 18 multi-omics data analysis tools in BnIR, including some common bioinformatics analysis tools, such as GO and KEGG enrichment analysis, BLAST, multi-sequence alignment (MSA), sequence extraction, variation annotation, primer design, electronic PCR and four graph visualization tools, which were all integrated into the Tools portal (Figure 3). Notably, most of analysis tools support 28 published *Brassica* genomes, and users can conduct these analyses without switching between different platforms. For population-level multi-omics data analysis, OnlineGWAS module can help users to perform GWAS by uploading phenotypic datasets. And we newly developed COLOC and SMR online analysis and visualization tools based on the rapeseed eQTL results of our database, which allow users to identify candidate genes and causal variants by uploading GWAS results. In addition, to facilitate the identification of germplasm resources, the SNPmatch tool based on the genetic variation dataset of 2,311 accessions was integrated into the Tool portal (Pisupati et al., 2017). In the SNPmatch module, the user can identify the accession with the highest similarity to their accession by uploading the genotype of the sample.

**Figure 3.**
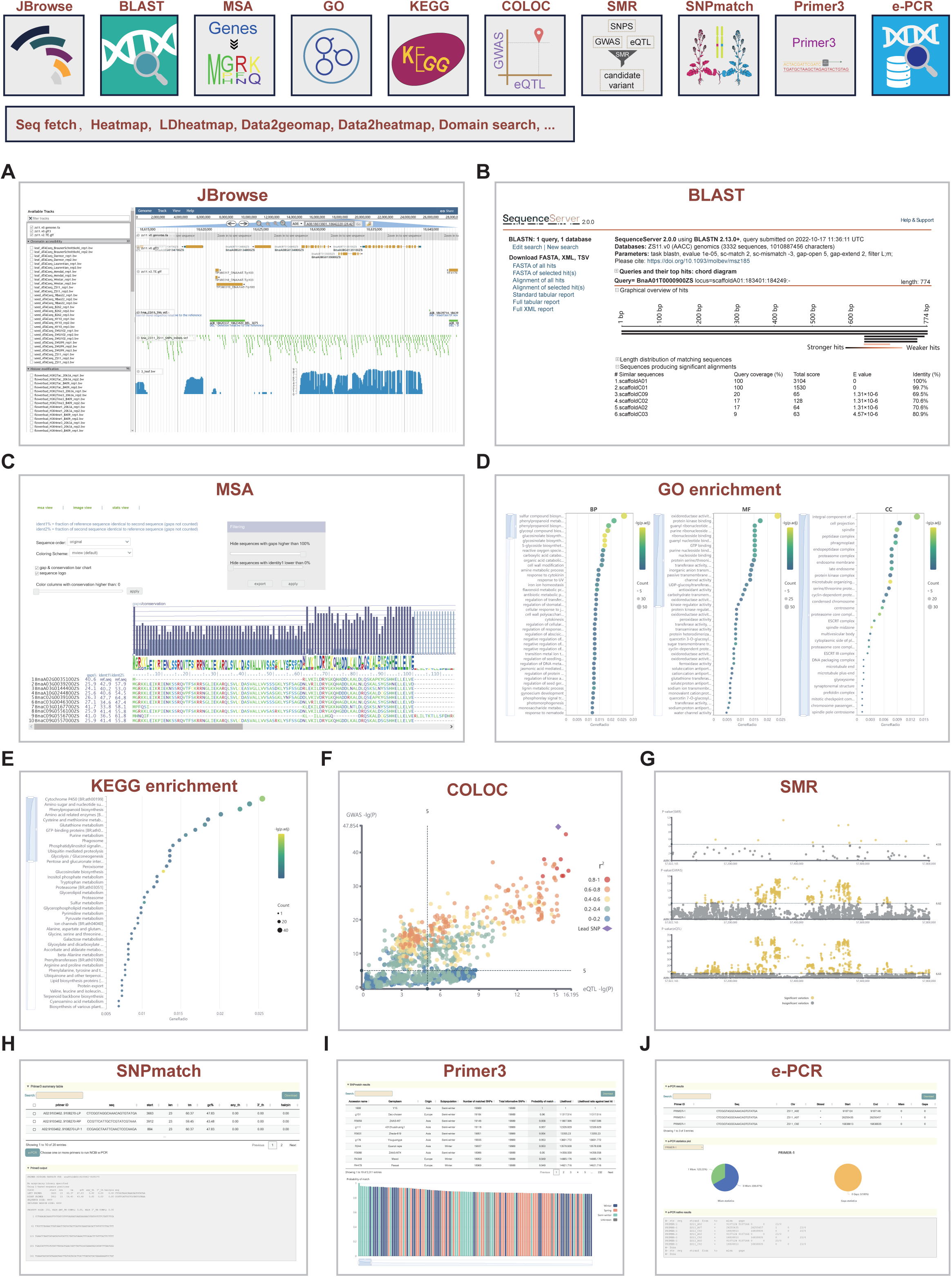
Multi-omics tools in the Tools portal of BnIR. A total of 18 tools were integrated into the Tools portal, including Jbrowse **(A)**, BLAST **(B)**, MSA **(C)**, GO enrichment analysis **(D)**, KEGG enrichment analysis **(E)**, co-localization analysis (COLOC) **(F)**, SMR **(G)**, germplasm identification tool SNPmatch **(H)**, primer design tool Primer3 **(I)** and electronic PCR tool e-PCR **(J)**.

In summary, BnIR provides abundant and comprehensive multi-omics data resources and convenient analysis tools. As a high-efficiency data analysis platform, BnIR allows direct utilization of these data and tools to mine genetic loci, genes and variations associated with specific traits. Furthermore, BnIR can also facilitate understanding of the mechanisms for gene expression regulation and phenotypic formation in the *B. napus* genome from different omics dimensions.

### Case study 1: Mining of genes associated with seed glucosinolate **content.**

SGC is an important trait related to the oil quality of rapeseed (Gupta and Pratap, 2007; Wang et al., 2018; Lu et al., 2019). In order to mine the genes related to SGC, GWAS was first performed. By using the GWAS module of the Multi-omics portal, the 8Glucosinolate content9 phenotype was selected for browsing significant GWAS signals (Figure 4A). As a result, a total of 20 genetic loci distributed in 10 chromosomes were identified to be associated with SGC (Figure 4B and Supplementary Table S5). Among them, *qGLS.C09.8* containing *BnaC09.MYB28a* and *BnaC09.MYB28b* was the most significant locus with the strongest effect (Figure 4C). *BnaC09.MYB28a* and *BnaC09.MYB28b* are members of the large family of R2R3-MYB transcription factors and have been identified as regulators of glucosinolate biosynthetic genes in *B. napus* (Harper et al., 2012; Wang et al., 2018). Notably, the most significant variations were distributed around *BnaC09.MYB28a* and *BnaC09.MYB28b* and showed strong linkage disequilibrium (LD) (Figure 4D). To further determine whether these two *BnaC09.MYB28s* are related to SGC, TWAS and co-localization analysis were performed. By using the TWAS module to browse the results, *BnaC09.MYB28a* and *BnaC09.MYB28b* were both found to be significantly associated with SGC (Supplementary Table S7), suggesting that their expression levels can potentially influence SGC. Then, the COLOC module was used to further test whether the GWAS locus and the *cis*-eQTLs of *BnaC09.MYB28a* and *BnaC09.MYB28b* are co-localized (Figure 4E). The two *BnaC09.MYB28s* and the 8Glucosinolate content9 phenotype were selected to query their co-localization analysis results (Figure 4E). As a result, they shared the same causal variant (PPH_4_ = 0.99), suggesting that *BnaC09.MYB28a* and *BnaC09.MYB28b* are candidate genes associated with SGC (Figure 4F and G). In addition, frameshift mutation (GCTA/-) may be the causal variation, which has been analyzed in our previous research (Wang *et al*., 2018; Yang *et al*., 2022a) (Figure 4F and G).

**Figure 4.**
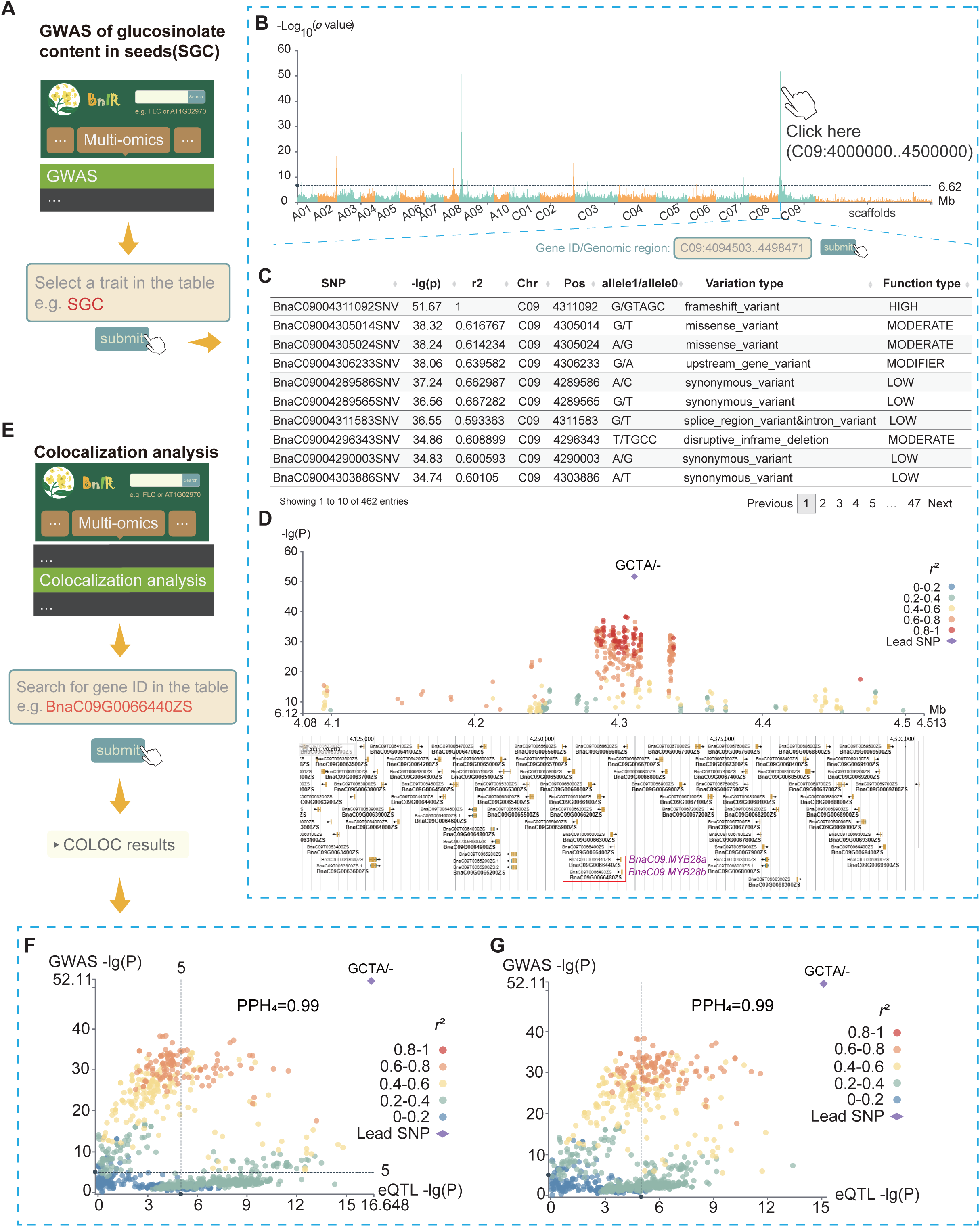
Case study of mining the genes associated with SGC. **(A)** Pipeline of searching SGC in GWAS module. **(B)** Genome-wide Manhattan plot of SGC in the GWAS module. **(C)** List of 402 significant variations associated with SGC identified by GWAS. **(D)** Local Manhattan plot of SGC at C09: 4.0–4.5 Mb. Two purple texts represent *BnaC09.MYB28a* and *BnaC09.MYB28b*. **(E)** Pipeline of searching the co-localization analysis results of *BnaC09.MYB28*s in BnIR. **(F-G)** Co-localization analysis of *cis*-eQTLs of two *BnaC09.MYB28*s and GWAS loci. *Cis*-eQTL signals of *BnaC09.MYB28a* **(F)** and *BnaC09.MYB28b* **(G)** co-localized with the *qGLS.C09.8* locus in GWAS. In **(D)**, **(F)** and **(G)**, purple diamonds represent the lead variations (one 4-bp frameshift InDel of GCTA/-), and green to red dots indicate weak to strong linkage disequilibrium coefficients with the lead variation.

### Case study 2: Identification of new genes associated with seed oil content

SOC is one of the most important traits of rapeseed (Wang *et al*., 2018; Tang *et al*., 2021). Here, the published SOC data of 286 *B. napus* accessions were used to validate the power of BnIR in identifying new candidate genes associated with traits (Wang *et al*., 2021). First, five significant loci were identified using GWAS, among which *qOC.C07.1* was the most significant locus. Then, the GWAS module in BnIR was used to browse the GWAS results in this locus (Supplementary Figure 1A). Significant variations were mainly distributed near *BnaC07.SRK2D* (BnaC07G0388200ZS) with strong LD (Supplementary Figure 1B). *BnaC07.SRK2D* encodes a member of abscisic acid (ABA)-activated SNF1-related protein kinases (*SnRK2*), which play important roles in various seed developmental processes such as de-greening, accumulation of seed storage products, seed maturation, desiccation tolerance and germination (Nakashima and Yamaguchi-Shinozaki, 2013; Lin et al., 2021; Ali et al., 2022; Kozaki and Aoyanagi, 2022). Next, we attempted to check whether the expression of *BnaC07.SRK2D* is related to SOC through co-localization analysis. When users enter the COLOC module of the Multi-omics portal and then select the eQTL of *BnaC07.SRK2D* and the QTL of SOC in GWAS (Figure 5A), the co-localization analysis results would be showed on the page (Figure 5B-D), which indicated that they shared the same causal variation, suggesting that *BnaC07.SRK2D* is a candidate gene for SOC.

**Figure 5.**
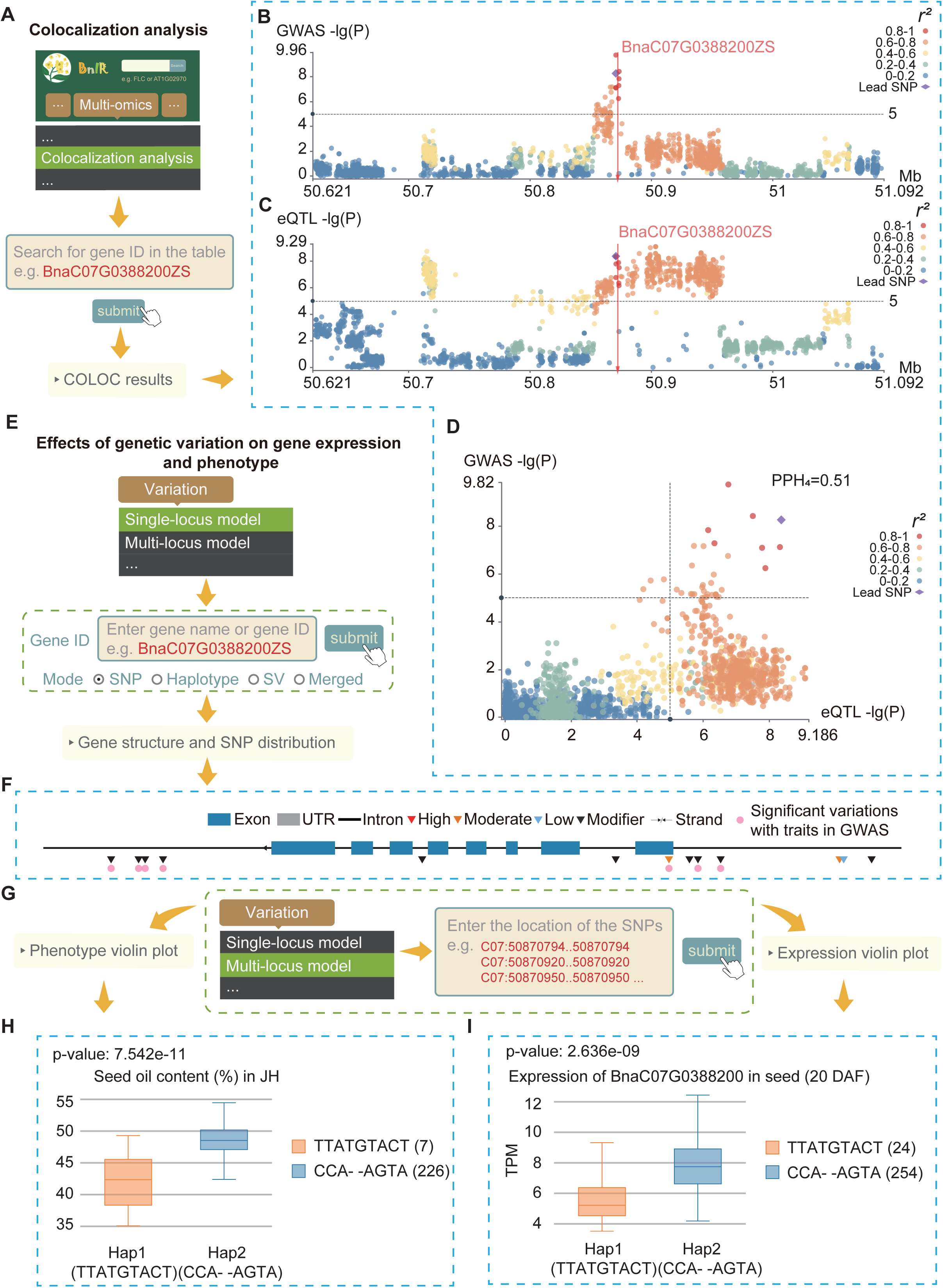
Case study of identifying new genes associated with SOC. **(A)** Pipeline of searching the co-localization analysis results of *BnaC07.SnRK2D* in BnIR. **(B-D)** Co- localization analysis of *cis*-eQTL of *BnaC07.SnRK2D* and GWAS loci. **(B-C)** Local Manhattan plots of SOC in GWAS **(B)** and *BnaC07.SnRK2D* in eQTL mapping **(C)**. **(D)** *Cis*-eQTL signals of *BnaC07.SnRK2D* co-localized with *qOC.C07.1* locus in GWAS. Purple diamond represents the lead variation. Green to red dots indicate weak to strong linkage disequilibrium coefficients with the lead variation. **(E)** Pipeline of searching variations in *BnaC07.SnRK2D* using the Single-locus module. **(F)** Variations in *BnaC07.SnRK2D* and its upstream and downstream variation within 1 kb. Red, orange, blue and black inverted triangles represent variations with high, moderate, low and modifier effects. Pink dots represent significant variations with SOC in GWAS. **(G)** Pipeline of haplotype analysis using the Multi-locus module. **(H-I)** Violin plots of SOC **(H)** and expression levels of *BnaC07.SnRK2D* **(I)** in accessions with Hap1 (orange boxes) and Hap2 (blue boxes).

Then, we used the Variation portal to further explore how genetic variation in this locus affects gene expression and ultimately influences the phenotype of the population. When entering 8BnaC07G0388200ZS9 in gene box of the Single-locus module of the Variation portal and choosing the 1 kb of flanking region (Figure 5E), we obtained 13 variations in this gene region, including ten variations in the upstream or downstream of the gene and three in the genic region (Figure 5F). Next, we performed a haplotype analysis to examine their effects on the gene expression and phenotype. When selecting these variations in the Multi-locus module (Figure 5G), we obtained two haplotypes with a frequency higher than 0.05 (Figure 5H and I). We found that the accessions with Hap2 (the same as Zhongshuang 11, a representative accession with high SOC; mean SOC, 48.51%) had significantly higher gene expression and SOC than those with Hap1 (mean SOC, 43.22%) (Figure 5H and I). These results indicated that these significant variations contribute to the divergence in the expression of *BnaC07.SRK2D* and phenotype. In other words, the accessions with Hap2 had significantly higher expression levels of *BnaC07.SRK2D*, which eventually contribute to higher SOC. Such findings will provide a valuable reference for increasing SOC in future breeding.

## Discussion

In this study, we first collected and processed rapeseed multi-omics data from recent research and databases, and then constructed BnIR, the first rapeseed multi-omics database. BnIR provides comprehensive multi-omics data resources, including genome sequences, annotations and expression levels of genes, epigenetic signals, metabolite contents, traits and association signals. In addition, multiple online analysis and visualization tools were developed and integrated to help researchers to mine genetic loci/genes and understand the regulatory mechanisms of gene expression and phenotypic formation. Two case studies including mining of candidate genes associated with SGC and SOC fully exhibited the power of BnIR in assisting genetic breeding.

Compared with the published rapeseed databases, BnIR has the following characteristics. 1) It integrates and optimizes the data sources and functions of BnPIR, BnTIR and BnVIR, and significantly improves the scale of the data source and response speed of the webpage (Figure 6A). 2) To the best of our knowledge, BnIR comprises the most comprehensive multi-omics datasets, including 13 published genome assemblies, genetic variations of 2,311 accessions, transcriptome data from 2,791 libraries, 118 phenotypes, contents of 266 metabolites, and multiple epigenome data involving histone modification, chromatin accessibility and chromatin interaction (Figure 6B). 3) GWAS, eQTL, TWAS, SMR and co-localization analysis are applied to mine the genetic loci, candidate genes and genetic variations, providing important resources, tools and references for rapeseed breeding (Figure 6B). 4) BnIR provides numerous common online multi-omics data analysis tools, which support all published rapeseed genomes and provide a convenient and efficient analysis platform.

**Figure 6.**
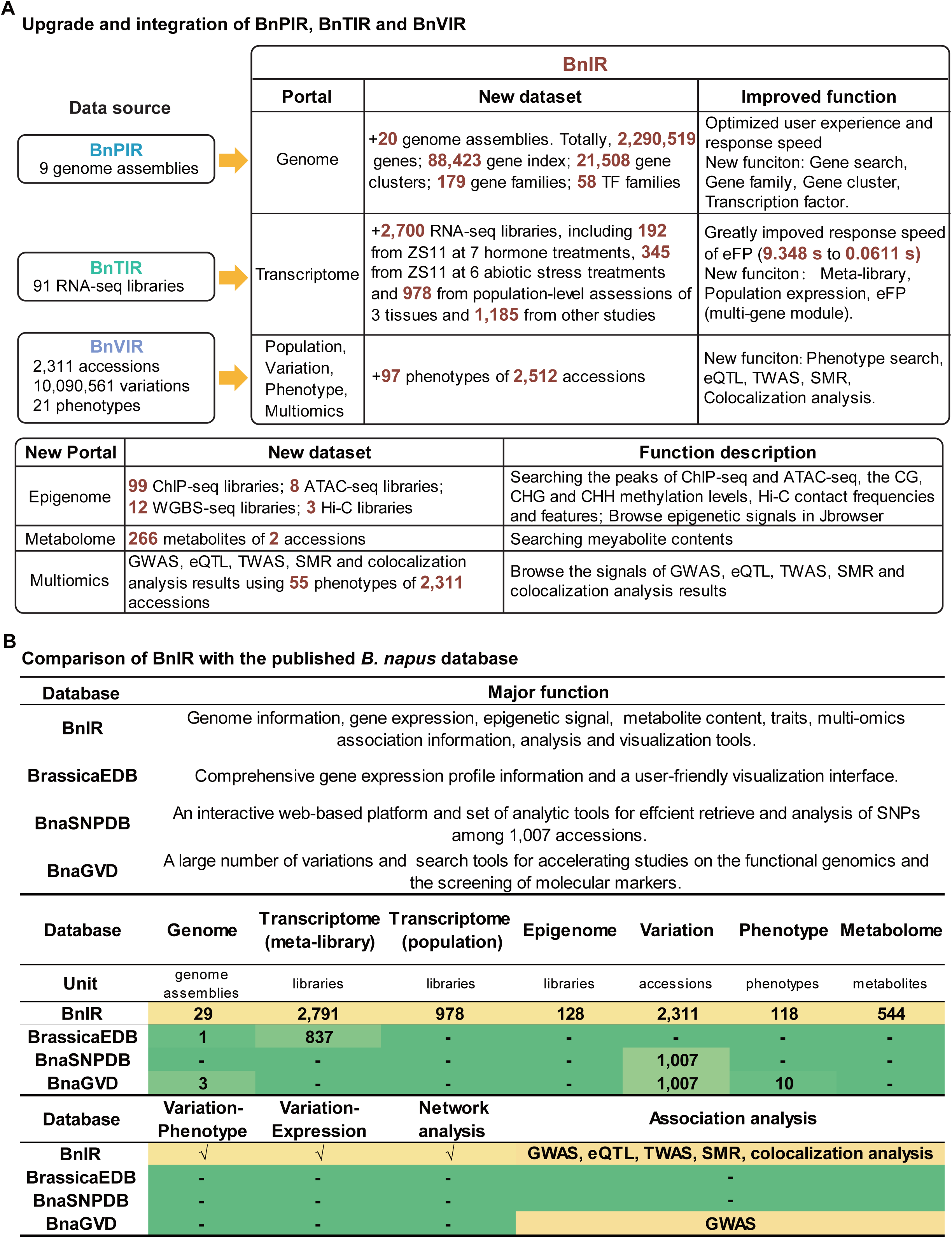
New features of BnIR. **(A)** Upgrade of BnIR relative to BnPIR, BnTIR and BnVIR. Red bold texts represent new datasets in BnIR. **(B)** Comparison of BnIR with other *B. napus* omics databases. Green to yellow cells represent fewer to more omics datasets.

In summary, we provide a valuable database for future rapeseed breeding and functional genomics research. In the future, with the release of more multi-omics datasets, we will update the database annually. In addition, more newly developed high-efficiency multi-omics methods will be applied to mine more candidate loci and genes associated with traits. These efforts will make BnIR a more useful platform in rapeseed research.

## Materials and Methods

### Data sources

To obtain comprehensive rapeseed multi-omics datasets, we mined and integrated the data from genomics, transcriptomics, genetic variations, phenotypic data, epigenetics and metabolomics (Figure 1; Supplementary Tables S1, S2, S3, S4). In total, 29 published *Brassica* genome assemblies (including 14 *B. napus* genome assemblies) and 2,290,519 genes in them were collected (Supplementary Table S1). As for the Transcriptome portal, 2,791 RNA-seq libraries of 272 individuals for 23 tissues were collected from published studies and databases (including 91 libraries from BnTIR) (Liu *et al*., 2021a), which are listed in Supplementary Table S2. As for the phenotypic data, the data of 118 phenotypes of 2,512 accessions were collected from 13 published studies (Figure 2; Supplementary Table S3). The epigenetics data are summarized in Supplementary Table S4. Metabolome datasets were retrieved from two previous studies (Geng et al., 2017; Clement et al., 2018). Genetic variation datasets of 2,311 accessions from BnVIR were also integrated into BnIR (Yang *et al*., 2022a).

### Reanalysis of omics datasets

For the omics datasets from different studies, considering the differences in data processing and analysis processes, unified pipelines and parameters were used for reanalysis of datasets during integration.

#### Genome annotation

For three newly collected *B. napus* genome assemblies, the repeat sequences were predicted as previously described and then integrated into the GBrowse of Genome synteny module (McKay et al., 2010; Song *et al*., 2020). The gene families, transcription factors, GO terms and KEGG pathways of all genome assemblies were predicted according to their homology to the *A. thaliana* genome and the annotations of *A. thaliana* downloaded from TAIR (https://www.arabidopsis.org/).

#### Comparative genome analysis

Genome sequences were compared between ZS11 reference genome and other 11 *B. napus* and two diploid (Z1 and HDEM) genome assemblies using the NUCmer program (v.4.0.0beta2) with parameters “nucmer--mum–noextend” in MUMmer4 (Marcais et al., 2018). After filtering of the one-to-one alignments with a minimum alignment length of 50 bp using the show-diff program from MUMmer4, the remaining alignment blocks were used for genome browser visualization. For genome browser visualization, the Dotplot module in JBrowse2 and Genome synteny module in GBrowser were embedded in BnIR (McKay *et al*., 2010; Hofmeister and Schmitz, 2018).

A total of 1,185,955 genes from 12 *B. napus* genome assemblies were used to construct the gene clusters. First, protein sequences of every pair from 12 genome assemblies were aligned using DIAMOND (v.0.9.14.115, http://github.com/bbuchfink/diamond). Then, gene synteny was detected by McScan (python version). The genes with synteny were grouped into one cluster. Finally, these genes were grouped into 231,106 gene clusters.

#### Transcriptome analysis

After clipping adaptor sequences and removing low-quality reads by fastp (v. 0.23.0), the clean RNA-seq data from accessions were mapped to the ZS11 reference genome using Hisat2 (v.2.1.2) with default parameters (Kim et al., 2015). Gene expression levels were normalized using the number of transcripts per kilobase million reads (TPM) by the StringTie software (v.1.3.5) with default settings(Pertea et al., 2015). The co-expression network was obtained by calculating the Pearson correlation coefficient of pairwise gene expression levels, and the gene modules including the gene pairs with a Pearson correlation coefficient higher than 0.8 were retained as a co-expression network.

#### Epigenome analysis

The methods used to remove adaptor sequences and filter low-quality reads were the same as those in RNA-seq data analysis. As for ChIP-seq and ATAC-seq, the clean data from accessions were mapped to the ZS11 reference genome using bowtie2 (v.2.3.2) with default parameters (Langmead and Salzberg, 2012). PCR duplicate reads were removed using Picard (v.2.19, http://broadinstitute.github.io/picard/). Peaks were called using the callpeak module of MACS2 software (v.2.1.2) with the parameters “--broad -f BAM -g 1000000000 -B -p 0.00001 --nomodel --extsize 147” (Feng et al., 2012).

As for Hi-C, the clean reads of each accession were mapped to the ZS11 reference genome using BWA-MEM with default parameters (Li and Durbin, 2009; Song *et al*., 2020). Then, the Hi-C interaction matrix was created using the Juicer pipeline (Durand et al., 2016). The KR-normalized matrix was extracted from Hi-C format files at the resolutions of 10 kb, 50 kb and 100 kb using the Juicer_tools (v.1.7.6) for JBrowser (Rao et al., 2014; Buels et al., 2016), and then compartments were called using Juicer_tools at the resolution of 100 kb.

As for the BS-seq data, clean data of each accession were mapped to the ZS11 reference genome using Bismark (v.0.13.0) with the parameter settings “-N 1, -L 30” (Krueger and Andrews, 2011). BigWig files of all epigenome data analysis can be visited by JBrowser in the Tools portal (Buels *et al*., 2016).

#### GWAS

As for the phenotypic datasets from different studies, GWAS was performed using the common pipeline. First, the SNPs and InDels with a minor allele frequency (MAF) lower than 0.05 were filtered. Then, GWAS was performed using GEMMA (v.0.98.1) (Zhou and Stephens, 2012). The population structure was controlled by including the first three principal components as covariates, as well as an IBS kinship matrix derived from all variants (SNPs and InDels) calculated by GEMMA. The cutoff for determining significant associations was set as -log_10_(1/n), where n represents the total number of variations. Finally, genetic loci and lead variation were identified with the following steps. 1) Two significant variations were regarded as the same locus if the distance between them was less than 0.5 Mb and; 2) As for a genetic locus, the most significant variation associated with the phenotype was identified as the lead SNP.

#### eQTL mapping

Gene expression values were taken as the values of the phenotype for eQTL mapping. Only those genes expressed in more than 95% of the accessions were defined as expressed genes for eQTL mapping. Variations with MAF > 5% were used to perform GWAS for each gene by using GEMMA to detect the associations for variations and genes (Zhou and Stephens, 2012). The cutoff for determining significant associations was set as -log_10_(1/n), where n represents the total number of variations. Based on the distance between eQTLs and targeted-genes, we subdivided all eQTLs into cis-eQTLs if the variation was found within 1 Mb of the transcription start site or transcription end site of the target gene, otherwise as trans-eQTLs. In BnIR, the regulatory pairs of eQTLs and eGenes are visualized using BioCircos.js (Cui et al., 2016).

#### TWAS

TWAS was used to integrate GWAS and gene expression datasets to identify gene-trait associations (Gusev *et al*., 2016). TWAS was conducted by the EMMAX module using the gene expression data of seeds at 20 DAF and 40 DAF with the data of the phenotypes from the same accessions. Models were considered as 8transcriptome-wide significant9 if they passed the Bonferroni correction for all genes.

#### SMR analysis

SMR analysis integrated the summary-level data from GWAS with eQTL data to identify genes associated with a complex trait considering their pleiotropy. The *cis*-eQTL signals of expressed genes and GWAS signals of the phenotype were used to perform SMR analysis and HEIDI test by SMR software (v.1.03) (Gusev *et al*., 2016). Then, the gene was defined as a candidate gene of the phenotype when the - log_10_(P-value) of SMR was lower than 1/n (n is the number of all expressed protein-coding genes) and P-value of HEIDI test was higher than 1.57 × 10^-3^.

#### Co-localization analysis

GWAS loci associated with phenotypes and candidate genes in these loci were used for co-localization analysis performed using the “COLOC” R package with default parameters (Hormozdiari et al., 2016). The variants in *cis*-eQTLs of genes and QTLs of phenotypes were defined as co-localized when the posterior probability of a co-localized signal (PPH_4_) value was higher than 0.5 and there was at least one shared significant variation.

#### GO and KEGG enrichment analysis

The GO and KEGG enrichment analysis was as follows: Firstly, we used blastp to establish homologous gene pairs of each rapeseed genome and Arabidopsis genome(Camacho et al., 2009; Boratyn et al., 2013), and then assigned the genes in the rapeseed genome according to the GO and KEGG annotation results of the Arabidopsis genome. Next, GO and KEGG libraries were established for each rapeseed genome using the 8clusterProfiler9 R package(Yu et al., 2012). Finally, based on the GO and KEGG libraries, the 8clusterProfiler9 R package was used for GO and KEGG enrichment analysis according to the gene list input by users.

### Implementation

BnIR (http://yanglab.hzau.edu.cn/BnIR) was constructed based on the ThinkPHP (v.5.0.24) framework with jQuery (v.3.6.0) as the JavaScript library, and runs on the Apache 2 web server (v.2.4.53) with MySQL (v.8.0.29) as its database engine. The database is available online without registration and optimized for Chrome (recommended), Opera, Firefox, Windows Edge and macOS Safari.

## DATA AVAILABILITY

Sources of all datasets are described in supplemental materials and methods. All datasets are made available at http://yanglab.hzau.edu.cn/BnIR/download.

## FUNDING

This research was supported by the National Natural Science Foundation of China (32070559), the National Key Research and Development Plan of China (2021YFF1000100), China Postdoctoral Science Foundation (2022M710875), Hubei Hongshan Laboratory (2021HSZD004) and Developing Bioinformatics Platform in Hainan Yazhou Bay Seed Lab (No. JBGS-B21HJ0001).

### Conflict of interest statement

None declared.

## Author contributions

Q.-Y.Y. designed the project. Q.-Y.Y. and Z.Y. managed and coordinated the project. Z.Y., S.W., Y.M.H., C.L., Y.J., Y.L. and Y.H. collected datasets and performed the bioinformatics analysis. Q.-Y.Y., Z.Y. and D.L designed the database; S.W., Y.M.H., L.W. and D.L. constructed the database. Q.-Y.Y., Z.Y., S.W. and Y.M.H wrote the manuscript. Q.-Y.Y., Y.Z., L.G. and C.D. directed the project.

## ACKNOWLEDGEMENTS

We thank Prof. Kede Liu from Huazhong Agricultural University of China, Prof. Kun Lu from Southwest University of China, Dr. Wujian from Yangzhou University of China, and other researchers for many very good and constructive feedback and suggestions in improving BnIR. We thank bioinformatics computing platform of the National Key Laboratory of Crop Genetic Improvement, Huazhong Agricultural University, managed by Hao Liu.

## Supplemental information

### Supplementary Figures

Supplementary Figure 1: Mining the genes associated with SOC using GWAS module.

### Supplementary Tables

Supplementary Table 1: Data source of 29 genome assemblies in the Genome portal.

Supplementary Table S2: Summary of transcriptome datasets in BnIR.

Supplementary Table S3: Summary of phenotypic datasets in BnIR.

Supplementary Table S4: Summary of epigenetics datasets in BnIR.

Supplementary Table S5: Significant GWAS loci associated with traits.

Supplementary Table S6: Summary of eQTL mapping.

Supplementary Table S7: Significant genes associated with traits by TWAS.

Supplementary Table S8: Significant genes associated with traits by SMR.

Supplementary Table S9: Significant genes associated with traits by co-localization analysis.

**Supplementary Figure 1:**
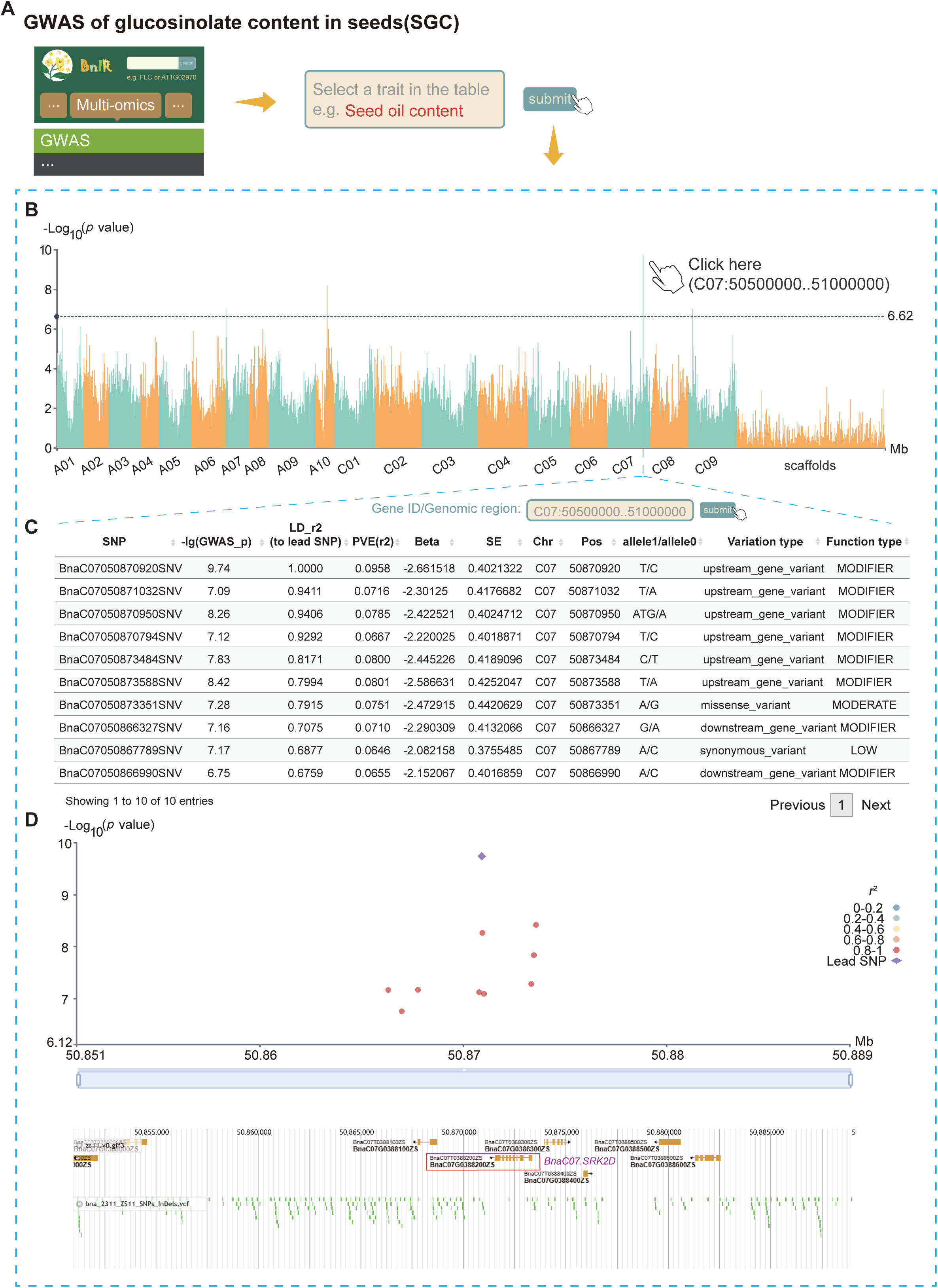
Mining the genes associated with SOC using GWAS module. **(A)** Pipeline of searching SOC in GWAS module. **(B)** Genome-wide Manhattan plot of SOC in the GWAS module. **(C)** List of 10 significant variations associated with SOC identified by GWAS. **(D)** Local Manhattan plot of SOC at C07: 50.5–51.0 Mb. The purple text represents *BnaC07.SRK2D*.

## Notes

http://yanglab.hzau.edu.cn/BnIR

### Competing Interest Statement

The authors have declared no competing interest.

### Summary of Updates

Manuscipt and Figures revised; author affiliations updated; Supplemental files updated.

